# Predicting mechanism of action of cellular perturbations with pathway activity signatures

**DOI:** 10.1101/705228

**Authors:** Yan Ren, Siva Sivaganesan, Nicholas A. Clark, Lixia Zhang, Jacek Biesiada, Wen Niu, David R. Plas, Mario Medvedovic

## Abstract

**Motivation:** Misregulation of signaling pathway activity is etiologic for many human diseases, and modulating activity of signaling pathways is often the preferred therapeutic strategy. Understanding the mechanism of action (MOA) of bioactive chemicals in terms of targeted signaling pathways is the essential first step in evaluating their therapeutic potential. Changes in signaling pathway activity are often not reflected in changes in expression of pathway genes which makes MOA inferences from transcriptional signatures a difficult problem.

**Results:** We developed a new computational method for implicating pathway targets of bioactive chemicals and other cellular perturbations by integrated analysis of pathway network topology, the LINCS transcriptional signatures of genetic perturbations of pathway genes and the transcriptional signature of the perturbation. Our methodology accurately predicts signaling pathways targeted by the perturbation when current pathway analysis approaches utilizing only a transcriptional signature of the perturbation fail.

**Availability and Implementation:** Open source R package paslincs is available at https://github.com/uc-bd2k/paslincs.

## INTRODUCTION

Misregulation of signaling pathway activity underlies many human diseases (Finkel and Gutkind, 2003; Laplante and Sabatini, 2012; Saxton and Sabatini, 2017). Identifying small molecules (i.e. chemical perturbagens) that can modulate activity of disease-related signaling pathways is the corner stone of intelligent drug design. This concept is exemplified by misregulation of the MTOR signaling pathways in various disorders and the activity of designing drugs to modulate MTOR signaling (Saxton and Sabatini, 2017). In the context of signaling pathways, the mechanism of action (MOA) of a biologically active molecule usually represents the direct effect that the molecule has on the activity of specific proteins in a pathway and therefore on the activity of the downstream elements within the pathway. The pathway MOA of bioactive molecules is important not only in assessing their therapeutic potential, but also their toxicity (Heijne, et al., 2005). In environmental toxicology, the target pathways are the essential component of the adverse outcome pathways framework aiming to predict the adverse health outcomes resulting from exposure to environmental exposures (Ankley, et al., 2010). The recently released dataset of perturbation transcriptional signatures (TS), consisting of genome-wide transcriptional changes after treatment with chemical perturbagens (CP)(Subramanian, et al., 2017), provides an opportunity to define MOAs of a large set of CPs. However, inferring the MOA from a transcriptional signature has been a difficult problem. The transcriptional signature represents a consequence of modulating signaling pathway activity while changes in activity of signaling proteins are often direct consequences of post-translational modifications and are not necessarily reflected in consistent changes in mRNA expression levels of corresponding genes (Dugourd and Saez-Rodriguez, 2019; Geistlinger, et al., 2016).

Nevertheless, there has been intense interest in inferring changes in the biological pathway activities based on the transcriptional signature (Khatri, et al., 2012; Mitrea, et al., 2013; Tarca, et al., 2013). Previous methods have ranged from simple statistical enrichment of differentially expressed genes among genes/proteins in the pathway (Tarca, et al., 2013) to network-based approaches attempting to assess consistency of the gene expression changes with the topology of protein-protein, protein-gene, and gene-gene interactions in the pathway(Mitrea, et al., 2013). Recent benchmarks of these and other methods have shown that the incorporation of pathway topology often yields very limited, if any, positive effect on the performance of different methods (Geistlinger, et al., 2016) which was attributed to the lack of changes in expression of pathway genes. On the other hand, Perturbation-RespOnse GENes (PROGENy) methodology used the transcriptional “footprints” of perturbed pathway genes, but without accounting for the topology of the pathway interaction network (Schubert, et al., 2018).

Gene expression changes after shRNA- or CRISPR-based knockdown of a gene can be used to precisely define a transcriptional signature of protein inactivation (Bild, et al., 2006). The concordance between such a genetic perturbation (GP) transcriptional signature and a transcriptional signature of a chemical perturbagen (CP), indicates the plausibility that the CP is perturbing the activity of the protein (Pilarczyk, et al., 2019; Subramanian, et al., 2017). In addition to CP signatures, recently released L1000 dataset generated by the LINCS project (Library of Integrated Network-based Cellular Signature) provides genetic perturbation signatures consisting of averaged changes in gene expression after knocking down the same gene with multiple shRNA’s for more than 3,500 human genes, perturbed in several cancer cell lines (Subramanian, et al., 2017). The new approach presented here leverages LINCS library of protein perturbation signatures to enable identification of the signaling pathways dysregulated by small molecules by integrated analysis of the CP signatures, the pathway network topology and GP signatures of pathway genes.

The key innovation of our methodology is the integration of two distinct strategies for implicating MOA of a chemical perturbagen: the topological pathway analysis (Mitrea, et al., 2013) and use of LINCS GP signatures (Pilarczyk, et al., 2019; Subramanian, et al., 2017). The integration is facilitated by an innovative statistical learning approach that uses the information about the topology of protein-protein interactions within a pathway and the LINCS GP signatures of the genes in the pathway to construct a pathway activity signature (PAS). We show that correlating transcriptional signatures of chemical perturbagens and other cellular perturbations with such PAS can implicate signaling pathways that are affected by the perturbation when standard methods fail to detect a signal. We also show how the new method can be used to refine pathway network models for specific biological contexts.

## Methods

### Transcriptional pathway activity signature (PAS)

Altered activity of a protein in a signaling pathway responding to chemical or genetic perturbation results in downstream changes in gene expression levels which are captured by the transcriptional signature. Our methods aim to identify the signaling pathways affected by the perturbation by comparing its transcriptional signature to GP signatures in the context of the pathway topology (Fig 1A). The key step in this process is the construction of the pathway activity signature (PAS) by integrating the topology of regulatory relationships within the pathway and the LINCS GP signaturees of genes in the pathway. Fig 1B-E illustrates the construction of the PAS on a small excerpt of the mTOR signaling pathway. The PAS is constructed in two steps: 1) *Signature genes* are selected by quantifying the consistency of changes in expression for each of the 978 measured L1000 landmark genes (Subramanian, et al., 2017) across the LINCS GP signatures of genes in the pathway with the pathway topology via a statistical model (Fig 1B-D); 2) The gene expression profiles of signature genes are summarized into a PAS (Fig 1E).

**Figure 1:**
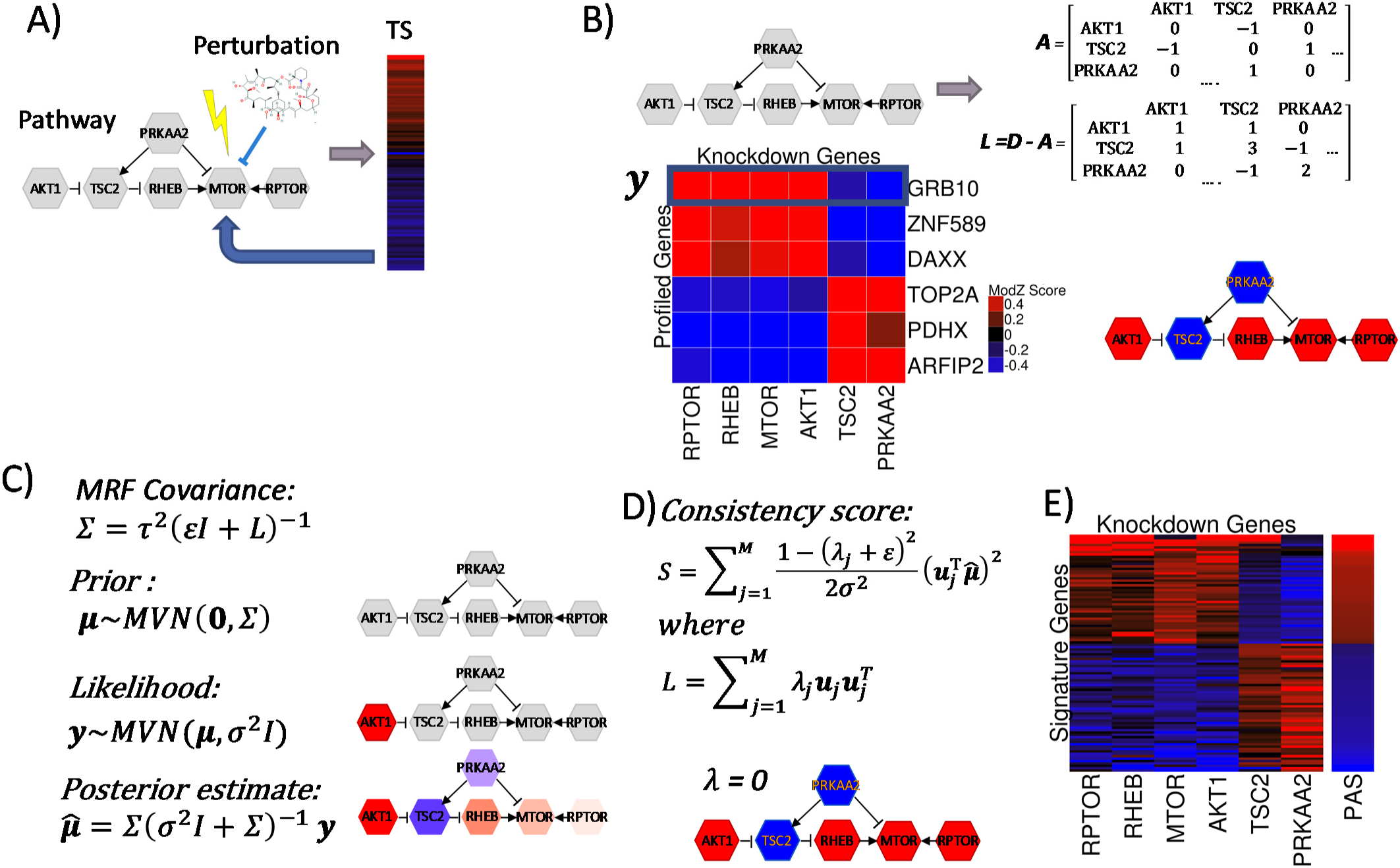
Integrating Signaling Pathway Topology and LINCS Consensus Gene Signatures to construct transcriptional Pathway Activity Signatures (PAS). In all panels, shades of red indicate different levels of positive and shades blue different levels of negative numbers. A) Chemical or genetic perturbation affects the activity of a protein in a signaling pathway and dysregulates the activity of the pathway. The pathway disregulation results in downstream changes in gene expression levels (gray arrow) which is captured by the transcriptional signature (TS). pasLINCS aims to identify the changes in pathway activity based on the downstream TS (blue arrow). B) The pathway activity signature (PAS) is constructed by integrating information from the LINCS genetic perturbation (GP) signatures of the pathway genes and the topology of protein-protein interactions in the pathway, summarized by the signed adjacency and Laplacian matrices. Expression profile of gene is consistent with the pathway topology if activation interaction between two nodes results in the expression change in the same direction and the inhibition interaction result in the change in the opposite direction C) Statistical model for integrating pathway topology with the expression profile of a gene; D) The gene-level consistency score between gene expression profile and the pathway topology. Top 100 genes with the highest consistency score are selected as “signature genes”; E) PAS consists of the gene loadings of the first principal component for the data matrix for signature genes across GP signatures of pathway genes.

To assess the consistency of the gene expression profile (***y***) of a single landmark gene across LINCS GP signatures of pathway genes with the pathway topology (Fig 1B), the pathway topology is summarized by the signed adjacency matrix *A* (Jacob, et al., 2012; Kunegis, et al., 2010). The assignment of positive (1) and negative (−1) weights to the edges in the pathway reflects the underlying assumption that GP signatures of two pathway genes should be positively correlated if the two genes are connected via “activating” interaction and negatively correlated if connected via inhibitory interaction. In the statistical model describing the distribution of the data (Fig 1C), the signed Laplacian (*L*) is then used as the precision matrix of the prior Markov random field for the mean expression changes (***μ***). The generative model for the data (***y***) is defined as the multivariate Gaussian distribution with mean ***μ*** and a diagonal variance matrix. Finally, the posterior mean vector 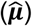 provides gene’s expected expression pattern after integrating the observed expression profile (***y***) and the pathway topology. The integration of the pathway topological structure and a gene expression profile by the statistical model is illustrated in Fig 1C for the gene expression profile representing the activation of a single node in the pathway (AKT1). The posterior mean estimate is consistent with the topology in the sense that its direction of the activity is consistent with the assumption that nodes connected by “activation” and “inhibition” relationships have positively and negatively correlated downstream effect on the gene expression changes respectively. Furthermore, the “closer” (in network topological sense) a node is to the initially activated node, the stronger is the activation signal. For each measured gene, we consider the norm of the projection of 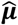 onto the lower dimensional subspace with the highest information content to represent the measure of consistency (ie *consistency score*) of its expression profile with the pathway topology (Fig 1D). Using the list of the 100 genes with highest consistency scores (*signature genes)*, PAS is constructed as the first principal component of the data matrix consisting of expression changes of signature genes in GP signatures of genes in the pathway (Fig 1D). A mathematical description of the statistical learning model is provided in the Supplementary Materials Section A.

A high-information subspace corresponds to a linear space spanned by the eigenvectors of the signed Laplacian corresponding to small eigenvalues, which have intuitive appeal from graph-theoretic perspective (Jacob, et al., 2012; Kunegis, et al., 2010). This can be shown by analyzing the Bayes factor (Kass and Raftery, 1995) for choosing between two probabilistic models that generated the data (***y***), one being the model in Fig 1C and the other being the model that assumes that topology of the pathway has no effect on the distribution of the data (Supplementary Materials Section B). We show that among the projections onto any one-dimensional subspace, the projection onto the null-subspace, the one corresponding to the eigenvector with the eigenvalue of zero, provides the highest discriminatory power for identifying genes with consistent expression profile (Supplementary Materials Section C). In our testing, going beyond the one-dimensional null-subspace did not improve significantly the discriminative ability of the signature (Supplementary Materials Section D). The eigenvector spanning the null space of the Laplacian (*λ* = 0), which is used to derive the gene activity scores, is visualized in Fig 1D. This illustrates the fact that basing the signature only on the projection to the null-space effectively summarizes the topology in terms of direction of the change in activity (increased, or decreased), but omits other aspects, such as distance of nodes in the network and the number of paths between different nodes that are captured by the rest of the eigenvectors.

### Node contribution score

For the purpose of assessing the contribution of individual GP signatures of pathway genes to the PAS, we use the *node contribution score.* The *node contribution score* is defined as the decrease in the consistency scores of signature genes after removing the GP signature of the node from the analysis. A positive node contribution score implies that the GP signature of the node improves the consistency of the expression profiles of signature genes with the pathway topology. Wilcoxon signed-rank test is used to test whether the node contribution is statistically significant.

### LINCS L1000 Consensus Gene Signatures and CP signatures

To construct GP and CP signatures used in the analyses, Level 4 LINCS L1000 dataset was downloaded from GEO (GSE92742). CP signatures were constructed by averaging Level 4 biological replicates. Level 5 moderated Z (MODZ) signatures of shRNA knock-downs were calculated as a weighted average of Level 4 replicates (Subramanian, et al., 2017). shRNA knock-down signatures were further integrated into GP signatures as weighted (MODZ) averages of individual shRNA signatures targeting the same gene. Only signatures designated as “gold” and generated using the “epsilon” version of L1000 probes were used in the analysis. All LINCS CP and GP signatures used in the analysis can be downloaded via *paslincs* package. The CP information of MOA is obtained from http://clue.io. Signatures of CPs that are activators or agonists of a target proteins were excluded from analyses.

### Baseline methods compared with PAS methodology

We considered six baseline methods to compare with our pasLINCS methodology. The baseline methods were designed to establish the benefits of including information from the GP signatures of pathway genes and the pathway network topology in the process of identifying targeted pathways. The first method (KD), defines a pathway signature as the first principal component of all GP signatures of the pathway genes. This method uses the GP signatures, but not utilizes the pathway network topology to identify informative genes. The second method (TP), regards a CP signature of the landmark genes in the pathway proteins as a gene profile, and calculates the consistency score for this profile. Then the consistency score is considered as a measure of the association between a pathway and a CP. This method is meant to represent the class of pathway analysis methodologies that utilize pathway topology to identify affected pathways based on the transcriptional data alone and does not use GP signatures of pathway genes. The third baseline method is the random set (RS) enrichment analysis, a prototypical pathway enrichment analysis method that does not make use of either pathway topology or GP signatures (Newton, et al., 2007). Additionally, the performance of three existing topological pathway analysis methods for analysis of transcriptional signatures was assessed: SPIA (Tarca, et al., 2009), CePa (Gu and Wang, 2013), and PathNet (Dutta, et al., 2012).

### ROC curves

For a specific target pathway, we focus on the TSes of CPs that inhibit any gene/protein within the pathway. For each of such TS, we designate all pathways not containing any protein/gene inhibited by the CP as true negatives, and calculate its false positive rate (FPR) as the proportion of correlations between the TS and PASes of true negative pathways that are larger than the correlation between the TS and the target pathway. For each FPR level, the corresponding true positive rate (TPR) is calculated as the proportion of all TSes targeting the pathway with FPR’s smaller than the given FPR level. ROC curves are then obtained by plotting FPRs against the corresponding TPRs. For each ROC curve, we calculate the area under the curve (AUC). We also calculated the partial area under the curve (pAUC) corresponding to the FPR<0.05 as this is a better measure of the precision of the methods in the relevant range of the specificity (Cheng, et al., 2014). We report the ratio of the pAUC to the area under the 45-degree line (rpAUC) as the measure of increase in the predictive ability over random predictions.

### Analysis of KEGG pathways

We processed *kgml* files corresponding to 328 homo sapiens KEGG pathways(Kanehisa, et al., 2017) using the R package *KEGGRest* to identify 179 “informative” pathways which contain at least two explicit “activation” or “inhibition” interactions and without “conflicting” interactions. For each informative pathway we constructed the signed adjacency matrix by setting the weights for “activation” edges to 1 and the weight for “inhibition” edges to −1, and calculating the signed Laplacian as shown in Fig 1. Supplementary Table S4 provides the summary of topological information contained in informative KEGG pathways. The pathways are grouped based on the secondary level classification in KEGG as: a.) pathways classified with the word “cancer” are grouped as “cancer”; b.) pathways classified related to a disease other than “cancer” are grouped as “disease”; c.) pathways classified with the word “signaling” or “signal” are grouped as “signaling”; d.) all other pathways are grouped as “other”

## RESULTS

### Transcription signature of mTOR signaling pathway activity

We studied the ability of our pasLINCS methodology to implicate genes whose expression pattern is a telltale sign of changes in pathway activity by constructing the PAS of mTOR pathway and by comparing it with transcriptional signatures of mTOR inhibitors. The protein interaction network representing mTOR signaling pathway was constructed by integrating information from KEGG and two recent papers describing the pathway (Saxton and Sabatini, 2017; Zhang, et al., 2017) (Fig 2A). The corresponding PAS showed a strong correlation with L1000 signatures of mTOR inhibitors in comparison with DMSO signatures (Fig 2B and 2C). To test whether the observed associations are platform independent we correlated PASes constructed from 12 LINCS cell lines with the time-course differential expression signatures of two glioma cell lines after treatment with a dual PI3K and mTOR inhibitor, PI-103 (Guillard, et al., 2009). Differential expression at 24 hours after PI-103 treatment in both glioma cell lines was significantly associated with the mTOR pathway PAS in the majority of the LINCS cell lines. Significant correlations can also be seen at 12 and 6 hours after treatment, but not before (Fig2D). These results are consistent with the expected dynamics of gene expression changes in response to PI-103 treatments (Guillard, et al., 2009). Similar analysis of the dataset studying the response of MCF-7 cell line to amino acid starvation (Tang, et al., 2015) showed consistent results with expected activation of mTOR signaling (Fig 2D). PASes constructed from three cell lines (NPC, SW480 and HCC515) showed lack of correlations in both analyses. SW480 cell line has previously been shown to be resistant to MTOR inhibition (Gulhati, et al., 2009), while the PAS for the NPC cell line was developed from only 6 CGSes. These factors along with relatively weak response (Fig 2E) to MTOR knockdown in HCC515 may explain the poor performance of PASes derived from these three cell lines.

**Figure 2:**
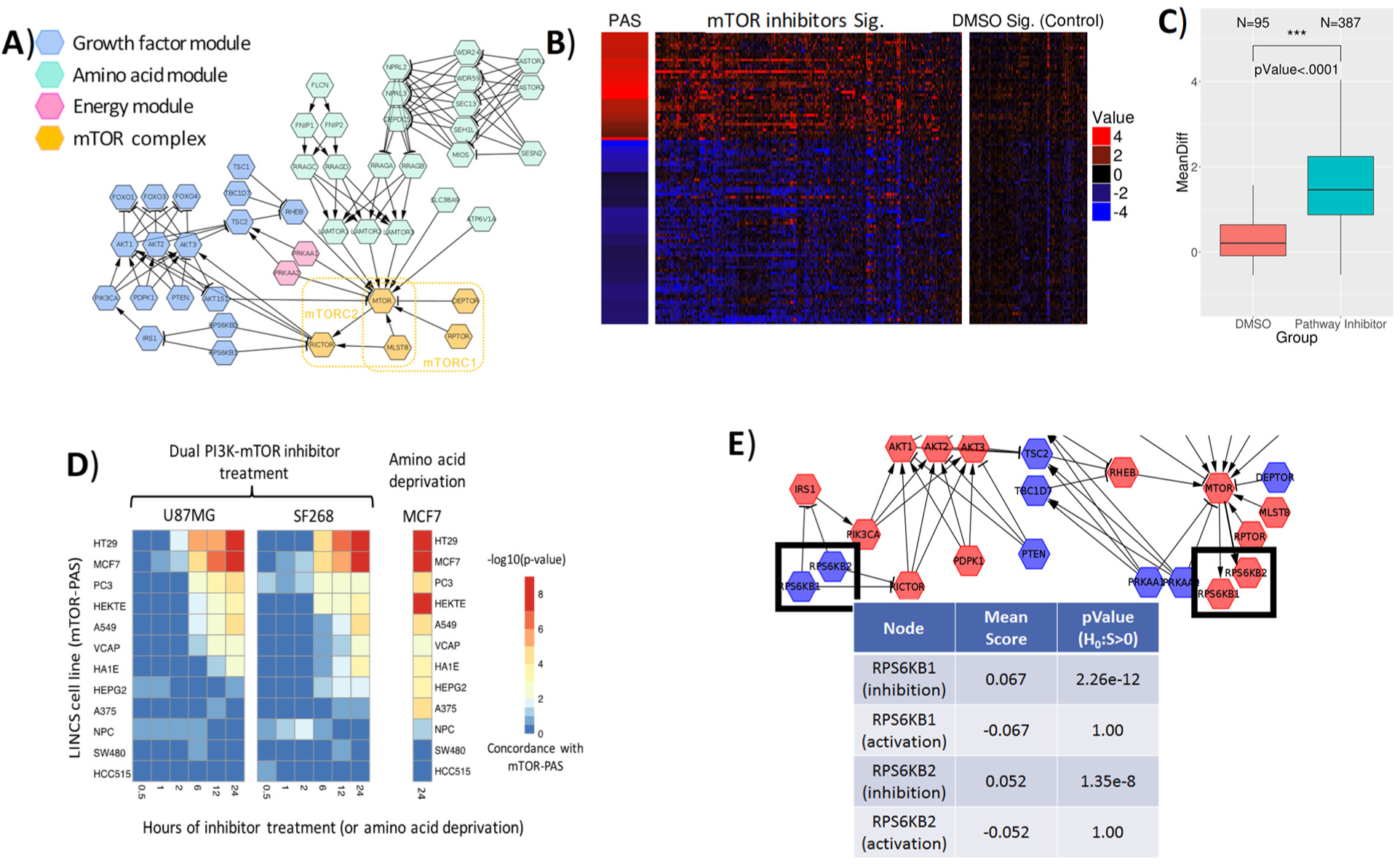
PAS of the mTOR signaling pathway. A) mTOR signaling pathway constructed from literature consisting of four key modules; B) PAS constructed using the methods in Fig 1 and the LINCS chemical perturbagen (CP) signatures for the pathway inhibitors and vehicle treatment; C) Distribution of differences in average expression levels between positive and negative signature genes for pathway inhibitors and vehicle treatment; D) Statistical significance of differences in expression level of positive and negative mTOR PAS genes after PI3K inhibition and amino acid starvation; E) Using node contribution scores to assess the role of S6K kinase in regulating mTOR pathway activity in the MCF7 PAS.

### Refining the pathway with node contribution scores

The pasLINCS can also be used to refine the pathway network by examining the changes in the pathway consistency scores for signature genes after removing one specific node. Nodes consistent with their implied role are expected to have a positive, statistically significant *node contribution score*. S6K proteins have been mapped as either upstream negative regulators or downstream positive output of mTORC1 activity in different biological contexts(Magnuson, et al., 2012; Shah and Hunter, 2006). Using node contribution scores, we studied the role played by S6K1 and S6K2 proteins in the upstream negative regulation (“feedback”) of mTORC1 and as downstream effectors of mTORC1 signaling. Genetic and biochemical data show that mTORC1 directly phosphorylates and activates S6K1 and on the other hand, S6K1 phosphorylates and destabilizes IRS1, which decouples upstream receptor tyrosine kinases from PI3K-mTORC1 signaling. These two roles result in conflicting positions in the pathway (Fig 2F), and, in any given context, the transcriptional signatures of their activity will be more consistent with only one of these roles. Using the node contribution scores for these two proteins under two topological models, we established that the expression signature of S6K protein knock-downs in L1000 data are consistent with their roles of inhibitors of mTOR signaling in the MCF7 cell line (Fig 2F) and the majority of other 8 cell lines (Supplementary Materials Section F).

### Predicting KEGG pathways affected by chemical perturbagens

We studied the ability of our methodology to identify KEGG signaling pathways modulated by a specific chemical perturbagen (CP). The evidence of CP effects on the activity of a pathway was assessed by the correlation between the CP transcriptional signature and the pathway’s PAS. For each KEGG pathway and for the our custom mTOR pathway, we constructed ROC curves evaluating the ability of such correlations to implicate pathways targeted by a CP. Fig 3A shows the ROC curve for the new method applied to mTOR signaling pathway (Fig 2A). For comparison, ROC curves are shown for methods that use information from only the LINCS GP signatures (KD), only the pathway topology (TP), and classical gene list enrichment that does not utilize either pathway topology or LINCS GP signatures (RS) (Fig 3A), as well as three topological pathway analysis methods (SPIA, PathNet and CePa). The ROC curves are summarized by the Area Under the Curve (AUC) and the area under the partial ROC curve (rpROC) for the high specificity (>0.9). Fig 3B shows the comparison of AUCs for all available KEGG pathways and in Fig3C detailed results for only signaling pathways are shown. The detailed information about performance for each pathway is shown in Fig S5 and Table S4, and summary results for rpAUC are shown in Fig S6. ROC analysis results indicate that: 1) pasLINCS methodology outperforms the methods based on simple enrichment analysis and topological pathway analysis methods that do not make use of GP signatures. The exact form of the topological pathway analysis method does not seem to make a big difference, with our simple and fast method (TP) overall performing on par with three state of the art topological pathway analysis methods.; 2) The use of the statistical model to identify signature genes based on the consistency of their expression profiles with the pathway topology improves the performance in comparison to using only GP signatures. 3) The use of GP signatures is particularly important in predicting affected signaling pathways with pasLINCS and KD methods outperforming methods that do not use GP signatures in 87% of signaling pathways

**Figure 3:**
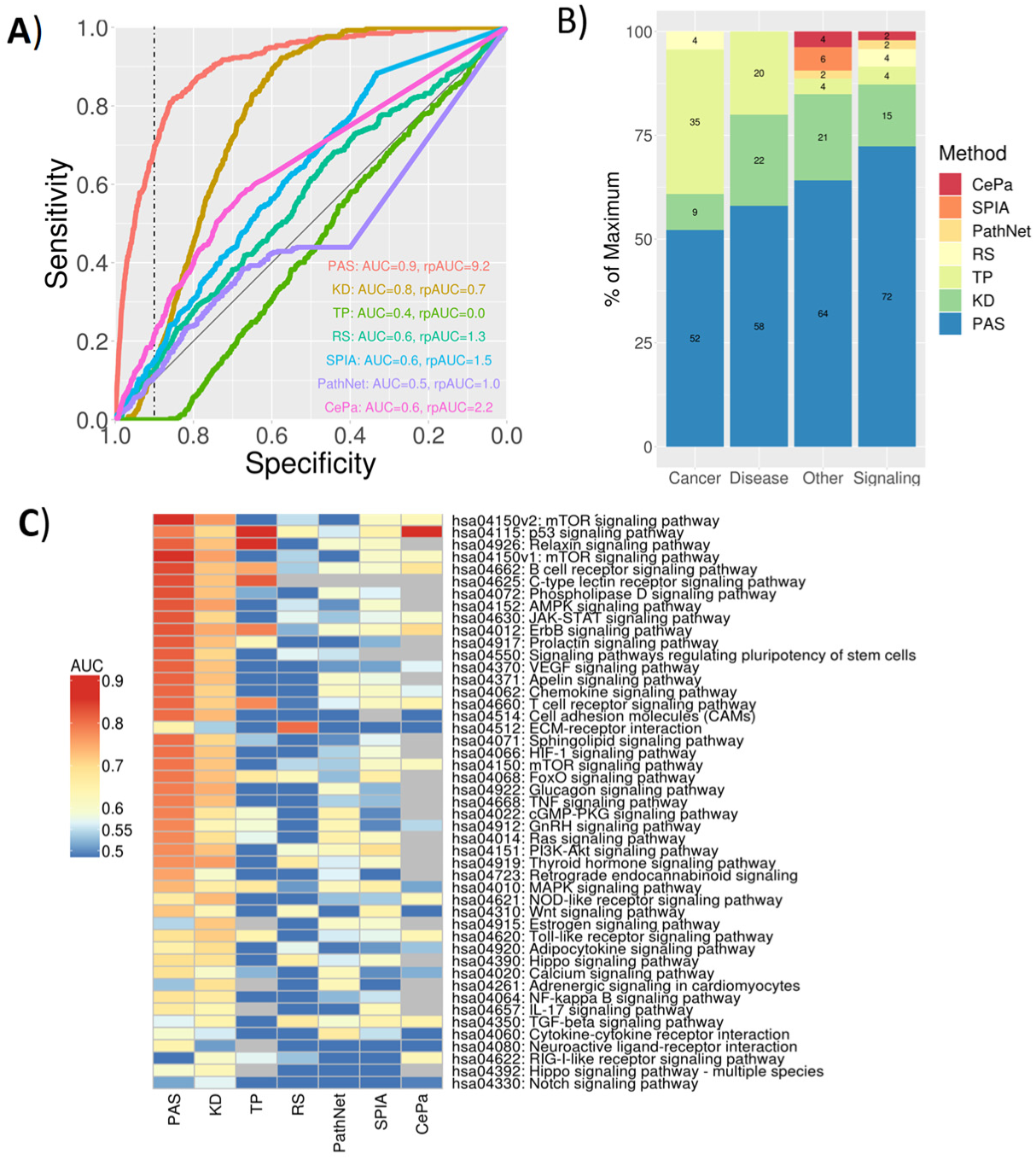
Predicting pathways perturbed by LINCS chemical perturbagens (CP). A) ROC curves for predicting correctly mTOR signaling pathways for CP’s known to target proteins in the pathway using seven different methods: PAS = pathway activity signatures using our new method; KD = pathway signatures constructed using only CGS data, but not utilizing the pathway topology; TP = using only CP transcriptional signatures and the pathway topology, but not using CGSes; RS = classical enrichment analysis not utilizing GP signatures or the pathway topology; B) Percentage of pathways predicted with the highest AUC for seven different methods across four different types of KEGG pathways; C) Heatmap of AUC’s for predicting affected KEGG signaling pathways for all seven methods.

## DISCUSSION

pasLINCS methodology integrates two distinct strategies for implicating signaling pathways affected by a CP based on its transcriptional signature: 1) the explicit modeling of shared expression changes implicated by the topology of the protein-protein interactions in the pathway (Mitrea, et al., 2013); and 2) Correlating CP transcriptional signature with the signatures of genetic perturbations of genes in the pathway (Pilarczyk, et al., 2019; Subramanian, et al., 2017). The use of GP signatures provides information about the activity changes in signaling proteins not contained in the CP transcriptional signature alone. Network based modeling integrates the information from different signaling proteins based on the expected interactions encoded by the pathway topology. Our results indicate that the new method is superior to either of the individual strategies in predicting signaling pathways affected by the bioactive chemical.

We compared the performance of our methods to three existing topological pathway methods (SPIA, PathNet and CePa) for analysis of transcriptional signatures that are applicable in analysis of LINCS CP signatures and do not require experimental replicates of treatment and control samples. They performed comparably with our own method (TP) and are likely representative of the results that can be obtained in general by a topological pathway analysis of CP signatures alone. In terms of methods that integrate multiple data types, PARADIGM uses a graphical model to integrate pathway network topology with Copy Number Variants (CNV) and gene expression profiles to identify pathway associated with cancer (Vaske, et al., 2010). Its graphical model, designed to integrate CNV and transcriptomic data, is not directly applicable for integration of GP and CP signatures.

Learning MOA of chemical perturbagens based on their transcriptional signatures opens new avenues for using connectivity map data to search for new therapies. In situations when the disease-related misregulation of signaling pathways is not clearly reflected in any available transcriptional signature, but is learned based on other information (e.g., genetics or proteomics studies), one can “connect” chemicals to disease based on their MOA. In the context of toxicogenomics, use of low-cost, high-throughput transcriptomic technologies (Bush, et al., 2017; Bushel, et al., 2018; Subramanian, et al., 2017), combined with pasLINCS analysis may open alternative avenues for high throughput safety evaluation of commercial chemicals, pesticides, food additives/contaminants, and medical products (Kavlock, et al., 2009; Kleinstreuer, et al., 2014). Previous studies have established the potential of assigning MOA of a chemical perturbagen based on comparison of their transcriptional signatures to the signatures of chemicals with known MOA (Iwata, et al., 2017; Subramanian, et al., 2017; Wang, et al., 2018). For example, the preciously derived PI3K inhibitor signature constructed from TSes of known chemical inhibitors (Zhang, et al., 2017) showed similar level of association with L1000 mTOR pathway inhibitors as we observed with our PAS (Fig 2D). pasLINCS adds another dimension by providing direct mechanistic link between the pathway activity and the effect of the CP without the need for reference signatures of perturbagens with known MOA.

The pasLINCS statistical learning model uses Bayesian inference to integrate the topological information with data on gene expression changes after perturbing nodes in the network. The key step in building the statistical model is the use of the regularized signed Laplacian as the precision matrix of the prior covariance to capture the effects of two basic kinds of protein-protein regulatory interactions in signaling cascades (activation and inhibition) on expression profiles of downstream genes. This simple representation is likely an oversimplification of the complexity of the dynamic biochemical processes taking place in transducing signals. However, our results show that the resulting covariance function captures adequately the static correlation structure of transcriptional signatures of pathway perturbations. The pasLINCS statistical learning model can be re-interpreted in the context of the regularization framework with graph kernels (Smola and Kondor, 2003) where standard Laplacian is replaced with the signed version. Similar strategies using standard graph with only positive edges have been used in the context of non-directed protein-protein interaction network (Cowen, et al., 2017) in general, as well as in predicting drug targets based on transcriptional signatures (Laenen, et al., 2013). The regularization formulation does not depend on the Gaussian distributional assumptions about the data used in our model, indicating that pasLINCS methodology is likely robust with respect to deviations from the distributional assumptions.

Our results demonstrate that pasLINCS methodology can be used to construct different variants of the pathway networks, but also postulate new hypotheses about the role that proteins may play in a signaling pathway. Analysis of the results indicated that the S6Ks GP signatures are more consistent with their role as inhibitors of the PI3K-AKT-mTOR signaling axes, but not as the transducers of mTORC1 activity. S6K1 is well established as a negative feedback regulator of insulin-stimulated AKT-mTORC1 signaling, while studies of S6K2 have revealed context-specific feedback function (Harrington, et al., 2004; Haruta, et al., 2000; Miller, et al., 2017; Pai, et al., 2016; Tremblay, et al., 2007). Optimization of node contribution scores led us to adopt the mTOR pathway network with both S6K1 and S6K2 inhibiting upstream pathway activation. Biologically, the observed results could also be explained by the fact that mTORC1 has multiple downstream transducers that affect transcriptional programs, in addition to S6K. Consequently, the inhibitory role of S6K affects more downstream transcriptional targets than its transducer role, which is then reflected in its GP signatures being dominated by its inhibitory role. Details of the S6K role may not be relevant to the goal of constructing an informative PAS, but it is easy to envision biological contexts in which such predictions would warrant reconfiguring the pathway with follow up experimentation to confirm the predicted role of a specific protein in the given context.

pasLINCS methodology opens a new avenue for functional analysis of transcriptomic data to discover mechanistic underpinnings of observed changes in gene expression levels. Our results indicate that in terms of implicating pathways affected by a CP, results of the pasLINCS analysis are complementary to the established enrichment strategies based on analysis CP transcriptional signatures alone. pasLINCS accurately predicts affected signaling pathways when established enrichment methods fail and should be included within general analytical pipelines for functional assessment of global changes in gene expression patterns.

## Supporting information

Supplementary Methods and Results

## FUNDING

This work has been supported by the National Institutes of Health [U54HL127624, P30ES006096].

## CONFLICT OF INTEREST

None

## Notes

#### Summary of Updates

This version has been edited to add comparisons to three existing topological pathway analysis methods.

