## Supplementary Methods and Results for "Predicting mechanism of action of cellular perturbations with pathway activity signatures"

### Supplementary Materials

##### *A. Statistical learning model for selecting signature genes and constructing PASes*

Suppose that for a single pathway with $M$ genes/proteins, $Y = (Y_{ij})$ is the $N \times M$ matrix of gene expression changes in $N$ ($N = 978$) measured landmark genes after genetically perturbing (i.e. knocking down) $M$ pathway genes. That is, the value $y_{ij}$ is the L1000 CGS measuring the differential expression level of the landmark gene $i = 1, \ldots, N$ after knocking down pathway gene $j = 1, \ldots, M$. The $i^{\text{th}}$ row in the matrix $Y$, $\boldsymbol{Y}_i = (Y_{i1}, Y_{i2}, \cdots, Y_{iM})^{\mathrm{T}}$ is the expression profile of the changes in expression in landmark gene $i$ across CGSes of pathway genes. Our statistical learning model is designed to discriminate between expression profiles of L1000 landmark genes that are consistent with the topology of the signaling pathway and those that are not.

Suppose that $A = (A_{ij})$ is the $M \times M$ signed adjacency matrix of the pathway network, that is

$$A_{ij} = \begin{cases} 1, & \text{if gene } i\ (j) \text{ activates gene } j\ (i); \\ -1, & \text{if gene } i\ (j) \text{ inhibits gene } j\ (i); \\ 0, & \text{if genes } i \text{ and } j \text{ are not connected.} \end{cases}$$

Let $L$ and $\Sigma$ be the signed Laplacian and the variance-covariance matrix of the Markov random field as defined in Fig 1. Furthermore, suppose that $Z_i = 1$ if gene $i$ is consistent with the pathway topology and $Z_i = 0$ otherwise. Then, we model the expression profile, $\boldsymbol{Y}_i$, of gene $i$ by $\boldsymbol{Y}_i | Z_i =$

$1 \sim MVN(\boldsymbol{\mu}_{i1}, \sigma^2 I)$ and $\boldsymbol{Y}_i | Z_i = 0 \sim MVN(\boldsymbol{\mu}_{i2}, \sigma^2 I)$, where $\boldsymbol{\mu}_{i1} \sim MVN(0, \Sigma)$ and $\boldsymbol{\mu}_{i2} \sim MVN(0, \tau^2 I)$. Suppose $\alpha$ is the probability for any one landmark gene to be consistent with the pathway, then the probability distribution for the profile of landmark gene $i$ is given by the two-component mixture of multivariate Gaussian distribution:

$$\boldsymbol{Y}_i \sim \alpha N(\boldsymbol{\mu}_{i1}, \sigma^2 I) + (1 - \alpha) N(\boldsymbol{\mu}_{i2}, \sigma^2 I).$$

The objective of our learning procedure is to evaluate the evidence in data ($\boldsymbol{Y}_i$) of $Z_i = 1$ vs. $Z_i = 0$. We achieve this by following the standard Bayesian learning approach and construct the consistency score based on the Bayes factor for comparing $Z_i = 1$ vs. $Z_i = 0$(Kass and Raftery, 1995). The consistency score is defined as

$$S_i = \sum_{j=1}^{M} \frac{1}{2\sigma^2} \left[1 - \left(\lambda_j + \varepsilon\right)^2\right] \left(\boldsymbol{u}_j^{\mathsf{T}} \hat{\boldsymbol{\mu}}\right)^2 \qquad \text{(Eq. S1)}$$

where $\hat{\boldsymbol{\mu}}$ is the Bayesian estimate of the mean expression profile assuming $Z_i = 1$ (Fig 1b), $\lambda_j$ and $\boldsymbol{u}_j$ are the $j^{\text{th}}$ ($j = 1,2,\cdots,M$) eigenvalue and the corresponding eigenvector of $L$.

The ratio $\sigma^2/\tau^2$ in our model defines the relative weight assigned to the pathway topology versus the expression data, and in the derivation of the score we set this ratio to 1. Since $\sigma^2$ is a constant multiplier in the consistency score for all genes, the exact value of $\sigma^2$ does not affect the comparison among genes (we set it to 1) (Eq. S1). To regularize the Laplacian we set $\varepsilon = 0.1$. However, in our analyses we use only the first eigenvector/eigenvalue ($k = 1$) and the exact value of $\varepsilon$ again does not affect the ranking of genes (Eq. S1). It can be shown (Supplementary Materials Section B) that $S_i$, which is a sum over all Laplacian eigen sub-spaces, is proportional to the logarithm of the Bayes factor for distinguishing $Z_i = 1$ from $Z_i = 0$. It can also be shown that, when $\varepsilon$ is small, the components of $S_i$ with the smallest eigenvalues are expected to contribute the most to the Bayes factor (Supplementary Materials Section C). The eigenvectors corresponding to small eigenvalues also capture the desirable properties of the expression profiles in terms of their consistency with the pathway topology (Jacob, et al., 2012). Therefore, we use only a subset of $k$ eigenvectors corresponding to the smallest $k$ eigenvalues. In our empirical tests (Supplementary Figure S2), the score based on only the smallest eigenvalue ($\lambda = 0$) performed very close to that with two smallest eigenvalues, as well as that with three smallest eigenvalues, and better than that with all eigenvalues.

For a pathway consisting of multiple connected subgraphs, we calculate the consistency score for each connected subgraph, and take the total of the scores as the final score to assess the consistency of the profiled gene to the pathway topology. Finally, we select $G$=100 of *signature genes* with the largest consistency scores corresponds to selecting $G$ genes with the highest posterior probabilities that their profiles are consistent with the pathway topology. In our tests (Supplementary Materials Section E) increasing the number of genes did not improve results. Using the knockdown signatures of the selected *signature genes*, we obtain the principal components of their covariance matrix, and use the eigenvector corresponding to the first principal component calculated as the pathway activity signature (PAS) (Fig 1D). The elements of a PAS corresponding to individual genes are referred to as the *activity scores*.

##### *B. Relating consistency score to the observed data and Bayes factor*

Suppose $\boldsymbol{Y} = (Y_1, Y_2, \cdots, Y_M)'$ is a gene profile. Let the first considered model $M_1$ defined as follows:

$$\boldsymbol{Y} \sim N(\boldsymbol{\mu}_1, \sigma^2 I), \text{ and } \mu_1 \sim N(\boldsymbol{0}, \Sigma),$$

where $\Sigma = \tau^2(L + \varepsilon I)^{-1}$ is a regularized inverse of the Laplacian $L$ of the pathway topology. Also let the second considered model $M_2$ defined as follows:

$$\boldsymbol{Y} \sim N(\boldsymbol{\mu}_2, \sigma^2 I), \text{ and } \mu_2 \sim N(\boldsymbol{0}, \tau^2 I).$$

Integrating out $\boldsymbol{\mu}_1$and $\boldsymbol{\mu}_2$, the marginal distributions of $\boldsymbol{Y}$ under these two models are:

$$\boldsymbol{Y} \sim N(\boldsymbol{0}, \sigma^2 I + \Sigma) \text{ and } \boldsymbol{Y} \sim N(\boldsymbol{0}, (\sigma^2 + \tau^2) I),$$

respectively.

Suppose $\Lambda$ is the diagonal matrix whose $i^{\text{th}}$ diagonal element $\lambda_i$ is the $i^{\text{th}}$ eigenvalue of $L$, and $U$ is the orthogonal matrix whose $i^{\text{th}}$ column $\boldsymbol{u}_i$ is the corresponding eigenvector of $L$. Thus we have the eigen-decomposition $L = U\Lambda U^T$, and with simple steps of algebra, we have

$$\sigma^2 I + \Sigma = \sigma^2 I + \tau^2 (U\Lambda U^T + \varepsilon I)^{-1} = UD_1U^T,$$

where $D_1 = Diag\left(\sigma^2 + \frac{\tau^2}{\lambda_1+\varepsilon}, \sigma^2 + \frac{\tau^2}{\lambda_2+\varepsilon}, \cdots, \sigma^2 + \frac{\tau^2}{\lambda_M+\varepsilon}\right)$.

By multiplying the likelihood function of the data ($\boldsymbol{Y}$) and the prior distribution of $\boldsymbol{\mu}$, it is derived that the posterior distribution of $\boldsymbol{\mu}$ is a normal distribution, and given a signature gene (which implies under $M_1$), the posterior mean expression profile is

$$\hat{\boldsymbol{\mu}} = \Sigma(\sigma^2 I + \Sigma)^{-1}\boldsymbol{Y} = UD_2U^T\boldsymbol{Y},$$

where $D_2$ is the diagonal matrix with the $i^{\text{th}}$ diagonal entry as

$$D_{2(ii)} = \frac{\tau^2}{\lambda_i+\varepsilon}\left(\sigma^2 + \frac{\tau^2}{\lambda_i+\varepsilon}\right)^{-1} = \frac{\tau^2}{\sigma^2(\lambda_i+\varepsilon)+\tau^2}.$$

Therefore, we have $U^T\hat{\boldsymbol{\mu}} = D_2U^T\boldsymbol{Y}$ and $U^T\boldsymbol{Y} = D_2^{-1}U^T\hat{\boldsymbol{\mu}}$.

On the other hand, the Bayes factor $B_{12}$ in favor of $M_1$ versus $M_2$ is given by

$$B_{12} = \frac{Likelihood(\boldsymbol{\mu}_1; \text{given } y)}{Likelihood(\boldsymbol{\mu}_2; \text{given } y)}$$

$$= \frac{|(\sigma^2 + \tau^2)I|}{|UD_1U^T|} \cdot exp\left\{-\frac{1}{2}(\boldsymbol{Y}^T U(D_1^{-1} - (\sigma^2 + \tau^2)^{-1} I)U^T \boldsymbol{Y})\right\}$$

$$= \prod_{i=1}^{M} \frac{\sigma^2+\tau^2}{\sigma^2+\frac{\tau^2}{\lambda_i+\varepsilon}} \cdot exp\left\{\frac{1}{2}(\boldsymbol{Y}^T UD_3U^T \boldsymbol{Y})\right\},$$

where $D_3$ is the diagonal matrix with the $i^{\text{th}}$ diagonal entry as

$$D_{3(ii)} = (\sigma^2 + \tau^2)^{-1} - \left(\sigma^2 + \frac{\tau^2}{\lambda_i+\varepsilon}\right)^{-1} = \frac{\tau^2(1-(\lambda_i+\varepsilon))}{(\sigma^2(\lambda_i+\varepsilon)+\tau^2)(\sigma^2+\tau^2)}.$$

We let $d = 1 + \frac{\sigma^2}{\tau^2}$. Without loss of generality, we can assume that $\tau^2 = \sigma^2$, and based on this assumption, we have $d = 2$. Then the diagonal matrix entries of $D_2$ can be simplified as $D_{2(ii)} = \frac{1}{(\lambda_i+\varepsilon)+1}$, and the exponential term of the Bayes factor $B_{12}$ can be further simplified with $D_{3(ii)} = \frac{1-(\lambda_i+\varepsilon)}{2\sigma^2((\lambda_i+\varepsilon)+1)}$.

Plugging in $U^T\boldsymbol{Y}$ to the exponential term of the Bayes factor $B_{12}$, we have

$$\boldsymbol{Y}^T UD_3U^T \boldsymbol{Y} = \widehat{\boldsymbol{\mu}}^T UD_2^{-1}D_3D_2^{-1}U^T \widehat{\boldsymbol{\mu}} = \widehat{\boldsymbol{\mu}}^T UD_4U^T \widehat{\boldsymbol{\mu}},$$

where $D_4$ is the diagonal matrix with the $i^{\text{th}}$ diagonal entry as

$$D_{4(ii)} = \left((\lambda_i + \varepsilon) + 1\right) \cdot \frac{1-(\lambda_i+\varepsilon)}{2\sigma^2((\lambda_i+\varepsilon)+1)} \cdot \left((\lambda_i + \varepsilon) + 1\right)$$

$$= \frac{1}{2\sigma^2}(1 - (\lambda_i + \varepsilon)^2).$$

Notice that the consistency score $S$ can be written in matrix form, and then related to the observed data by substituting $\hat{\mu}$ by $Y$ as follows:

$$S = \sum_{j=1}^{M} \frac{1}{2\sigma^2}\left[1 - (\lambda_j + \varepsilon)^2\right](\boldsymbol{u}_j^T \widehat{\boldsymbol{\mu}})^2$$

$$= \widehat{\boldsymbol{\mu}}^{\boldsymbol{T}} UD_4U^T \widehat{\boldsymbol{\mu}}$$

$$= \boldsymbol{Y}^T UD_3U^T \boldsymbol{Y}.$$

Because of the relationship between the data ($\boldsymbol{Y}$) and the estimate of the mean expression profile ($\widehat{\boldsymbol{\mu}}$), we can easily calculate the consistency score by using the eigen-components and data without estimating the posterior mean.

Furthermore, the Bayes factor $B_{12}$ is associated to the consistency score $S$ as

$$B_{12} = \prod_{i=1}^{M} \frac{2(\lambda_i + \varepsilon)}{\lambda_i + \varepsilon + 1} \cdot exp\left\{\frac{1}{2}\widehat{\boldsymbol{\mu}}^{\boldsymbol{T}} U D_4 U^T \widehat{\boldsymbol{\mu}}\right\}$$

$$= \prod_{i=1}^{M} \frac{2(\lambda_i+\varepsilon)}{\lambda_i+\varepsilon+1} \cdot exp\left\{\frac{1}{2} S\right\}.$$

This relationship shows that the consistency score is twice of the exponential term of the Bayes factor, and it motivates us to define the consistency score as such to detect the consistency of a gene to a pathway topology. Note that when we use only the smallest eigenvalue ($\lambda_1 = 0$) and small value of ($\varepsilon \ll 1$), the corresponding diagonal entry of $D_3$ (and that of $D_4$) is a positive constant, and the larger value $\boldsymbol{Y}^T \boldsymbol{u}_1 \boldsymbol{u}_1^T \boldsymbol{Y}$ (as well as the value $\widehat{\boldsymbol{\mu}}^T \boldsymbol{u}_1 \boldsymbol{u}_1^T \widehat{\boldsymbol{\mu}}$ ) corresponds to the larger Bayes factor $B_{12}$ for a specific pathway topology. This motivates us to select the signature genes with large consistency scores.

##### *C. Motivation for the use of projection to the null-subspace – comparison to other single subspace*

Suppose $\boldsymbol{Y} = (Y_1, Y_2, \cdots, Y_M)'$ is a gene profile. Let the first considered model $M_1$ defined as follows:

$$\boldsymbol{Y} \sim N(\boldsymbol{\mu}_1, \sigma^2 I), \text{ and } \mu_1 \sim N(\boldsymbol{0}, \Sigma),$$

where $\Sigma = \tau^2 (L + \varepsilon I)^{-1}$ is a regularized inverse of the Laplacian $L$ of the pathway topology. Also let the second considered model $M_2$ defined as follows:

$$\boldsymbol{Y} \sim N(\boldsymbol{\mu}_2, \sigma^2 I), \text{ and } \mu_2 \sim N(\boldsymbol{0}, \tau^2 I).$$

Integrating out $\boldsymbol{\mu}_1$ and $\boldsymbol{\mu}_2$, the marginal distributions of $\boldsymbol{Y}$ under these two models are:

$$\boldsymbol{Y} \sim N(\boldsymbol{0}, \sigma^2 I + \Sigma) \text{ and } \boldsymbol{Y} \sim N(\boldsymbol{0}, (\sigma^2 + \tau^2) I),$$

respectively. Under $M_1$, the profile $\boldsymbol{Y}$ is concordant to the pathway topology, while under $M_2$ the profile is completely random and unrelated to the pathway topology.

Let $\widetilde{\boldsymbol{Y}} = U^T\boldsymbol{Y}$, where $U$ is the orthogonal matrix, so that $U^TLU = Diag(\lambda_1, \lambda_2, \cdots, \lambda_M)$, where $\lambda_1, \lambda_2, \cdots, \lambda_M$ are the eigenvalues of $L$. Then the marginal likelihood function under models $M_1$ and $M_2$ are given, respectively, by

$$L_1 = f_N(\widetilde{\boldsymbol{Y}}: \boldsymbol{0}, D_1), \text{ and } L_2 = f_N(\widetilde{\boldsymbol{Y}}: \boldsymbol{0}, (\sigma^2 + \tau^2)I),$$

where $D_1 = Diag(\sigma^2 + \frac{\tau^2}{\lambda_1+\varepsilon}, \sigma^2 + \frac{\tau^2}{\lambda_2+\varepsilon}, \cdots, \sigma^2 + \frac{\tau^2}{\lambda_M+\varepsilon})$, and $f_N(\cdot)$ is the multivariate normal density.

The logarithm of Bayes factor $B_{12}$ in favor of $M_1$ versus $M_2$ is given by

$$2logB_{12} = log\frac{|(\sigma^2 + \tau^2)I|}{|UD_1U^T|} - \widetilde{\boldsymbol{Y}}^T(D_1^{-1} - (\sigma^2 + \tau^2)^{-1}I)\widetilde{\boldsymbol{Y}}$$

$$= \sum_{i=1}^{M}\left(log\frac{\sigma^2+\tau^2}{\sigma^2+\frac{\tau^2}{\lambda_i+\varepsilon}} - \frac{\tau^2(\lambda_i+\varepsilon-1)}{(\sigma^2(\lambda_i+\varepsilon)+\tau^2)(\sigma^2+\tau^2)}\tilde{y}_i^2\right).$$

Large positive logarithm of the Bayes factor $B_{12}$ would indicate that the data is more favorable in the model $M_1$. The expectation of the logarithm of Bayes factor $B_{12}$ under $M_1$ is given as follows:

$$E(2logB_{12}) = \sum_{i=1}^{M}\left(log\frac{\sigma^2+\tau^2}{\sigma^2+\frac{\tau^2}{\lambda_i+\varepsilon}} - \frac{\tau^2(\lambda_i+\varepsilon-1)}{(\sigma^2(\lambda_i+\varepsilon)+\tau^2)(\sigma^2+\tau^2)}\cdot\left(\sigma^2 + \frac{\tau^2}{\lambda_i+\varepsilon}\right)\right).$$

We let $d = 1 + \frac{\sigma^2}{\tau^2}$, and $g(\lambda_i) = \frac{1}{\lambda_i+\varepsilon} - 1$. Then we have

$$E(2logB_{12}) = \sum_{i=1}^{M}\left(log\frac{d}{d+g(\lambda_i)} + \frac{g(\lambda_i)}{d}\right) \stackrel{\text{def}}{=} \sum_{i=1}^{M}h(\lambda_i).$$

The contribution of the $i^{th}$ term $h(\lambda_i)$, which corresponds to the $i^{th}$ eigen component, is the contribution to the expected logarithm of Bayes factor $B_{12}$, and is indicative of its discriminatory power when $M_1$ is true. The larger contribution is, the more discriminatory power this term has, under $M_1$.

Without loss of generality, we can assume that $\tau^2 = \sigma^2$, and based on this assumption, we have $d = 2$, and $h(\lambda) = log\frac{2}{2+g(\lambda)} + \frac{g(\lambda)}{2}$. And when $0 < \varepsilon \ll 1$, we have

$$h(0) = log\left(\frac{2\varepsilon}{1+\varepsilon}\right) - \frac{\varepsilon - 1}{2\varepsilon} > 0$$

$$h'(\lambda) = \frac{\varepsilon + \lambda - 1}{2(\varepsilon+\lambda)^2(\varepsilon+\lambda+1)} \begin{cases} < 0, & \text{when } \lambda<1-\varepsilon \\ \geq 0, & \text{when } \lambda \geq 1-\varepsilon \end{cases}$$

and $\lim_{\lambda\to\infty} h(\lambda) = log2 - \frac{1}{2}$.

Hence we know that for small value of $\varepsilon$ which satisfies $h(0) > \lim_{\lambda\to\infty} h(\lambda)$, the function $h(\lambda)$ decreases as $\lambda$ increases from 0 to 1-ε, and then increases to the upper bound as $\lambda$ increases from 1-ε to infinity. This indicates that the highest discriminatory power is achieved when λ=0. We plot $h(\lambda_i)$ vs. $\lambda_i$ with four different small values of ε to show the contribution of different components corresponding to the different eigen values $\lambda_i$ (Figure S1).

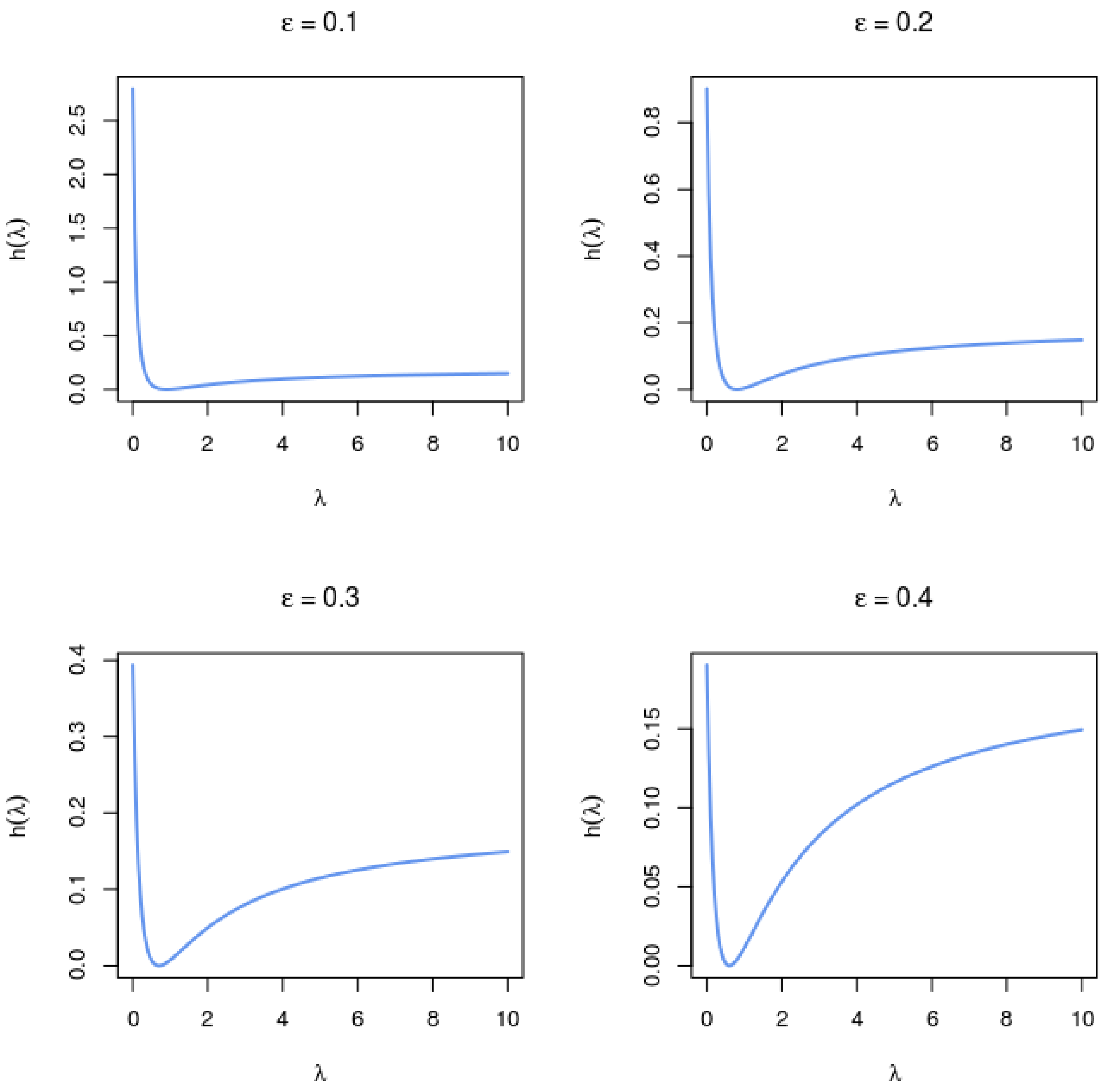

Figure S1. $h(\lambda)$ versus $\lambda$ with different values of $\varepsilon$

R code for plotting Figure S1:

```
par(mfrow=c(2,2))

l=seq(0,10,0.05)

h <- function(lambda,epsilon) {

          g <- 1/(lambda+epsilon)-1

          return(log(2/(2+g))+g/2)

     }

plot(l,h(l,0.1),xlab=expression(lambda),ylab=expression(h(lambda)),type="l",lwd=2,col="cornflowerblue",main=expression(paste(epsilon," = 0.1")))

plot(l,h(l,0.2),xlab=expression(lambda),ylab=expression(h(lambda)),type="l",lwd=2,col="cornflowerblue",main=expression(paste(epsilon," = 0.2")))

plot(l,h(l,0.3),xlab=expression(lambda),ylab=expression(h(lambda)),type="l",lwd=2,col="cornflowerblue",main=expression(paste(epsilon," = 0.3")))

plot(l,h(l,0.4),xlab=expression(lambda),ylab=expression(h(lambda)),type="l",lwd=2,col="cornflowerblue",main=expression(paste(epsilon," = 0.4")))
```

##### *D. Motivation for the use of projection to the null-subspace – comparison to higher dimensional subspaces*

We empirically compared the performance of consistency scores between using only the projection on the null-subspace ($\lambda = 0$) and using the projections on multiple subspaces. We considered using the smallest eigenvalue ($\lambda = 0$) as the baseline, and compared to the results of using the smallest 2 and 3 eigenvalues, and all eigenvalues. Our comparison is based on the TSes of MCF7 cell line. According to Figure S1, the smallest several values of $\lambda$ dominate more when $\varepsilon = 0.1$. Based on the L1000 signature data cleaning steps, it is reasonable to assume the unit variance for the samples ($\sigma^2 = 1$). Here we compared the performance of PAS through the ROC curves of benchmarking 173 pathways using the TSes of CPs based on $\varepsilon = 0.1$. Table S1 and Figure S2 show the statistics and distributions of AUC and rpAUC using two, three, and all eigenvalues compared to using only one smallest eigenvalue. Using the second smallest eigen-component provides slightly improvement in the performance. This may be because the second eigenvector indicates the “distance” of nodes in the topology. No improvement in the performance of PAS was observed by using three eigen-components corresponding to the smallest three eigenvalue, and using all eigen-components even performed slightly worse than only projecting onto the null subspace. Hence for simplicity and performance, we suggest users to use only the smallest one or two eigenvalues to calculate the consistency score.

|  | N_Eigen | Diff_Mean | Diff_Median | p.value |
|---|---|---|---|---|
| AUC | Two | 0.003 | 0.002 | 0.0264 |
|  | Three | -0.001 | 0.000 | 0.5079 |
|  | All | -0.069 | -0.071 | 1.0000 |
| rpAUC | Two | 0.070 | 0.035 | 0.0123 |
|  | Three | -0.047 | 0.000 | 0.4366 |
|  | All | -0.327 | -0.191 | 0.9961 |

Table S1. Statistics of differences in AUC and rpAUC. The differences are calculated as the corresponding measurement obtained by using the additional eigenvalues (2, 3, and all) subtracted by the measurement obtained by using only the smallest eigenvalue $\lambda = 0$. The third column is the average of differences for 173 pathways. The fourth column is the median of differences for 173 pathways. The last column indicates the p-value of the one-sided (greater than zero) non-parametric Wilcoxon signed rank test comparing the difference and zero.

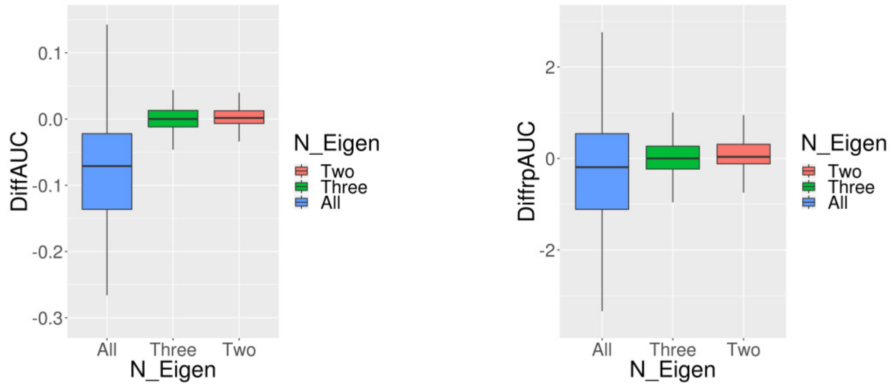

(a) (b)

Figure S2. Distribution of differences in AUC and rpAUC of using the smallest up to three and all eigenvalues compared with using only the smallest eigenvalue when calculating consistency scores. (a) Difference in AUC. (b) Difference in rpAUC.

#### *E. Sensitivity of the number of signature genes*

We assessed the sensitivity of the number of signature genes by comparing the performance of PAS through the ROC curves of benchmarking 173 pathways based on MCF7 cell line with 50, 100, 150, and 200 signature genes. Table S2 and Figure S3 show that the performance of using 100 signature genes is not worse than that of using 50 or 200 signature genes with respect to AUC and rpAUCs, and equally good to that of using 150 signature genes with respect to AUC and slightly lower in rpAUC. This implies that the number of top genes used as signature genes within the range of 50 to 200 is not sensitive to the performance.

| | N_Top | Diff_Mean | Diff_Median | p.value |
|---|---|---|---|---|
| AUC | 50 | -0.004 | -0.007 | 0.9995 |
| | 150 | -0.001 | -0.001 | 0.4632 |
| | 200 | -0.006 | -0.005 | 0.9824 |
| rpAUC | 50 | -0.187 | -0.023 | 0.9949 |
| | 150 | 0.089 | 0.000 | 0.0346 |
| | 200 | 0.088 | 0.000 | 0.1418 |

Table S2. Statistics of differences in AUC and rpAUC. The differences are calculated as the corresponding measurement obtained by using 50, 150, and 200 signature genes subtracted by the measurement obtained by using 100 signature genes. The third column is the average of differences for 168 pathways. The fourth column is the median of differences for 168 pathways. The last column indicates the p-value of the one-sided (greater than zero) non-parametric Wilcoxon signed rank test comparing the difference and zero.

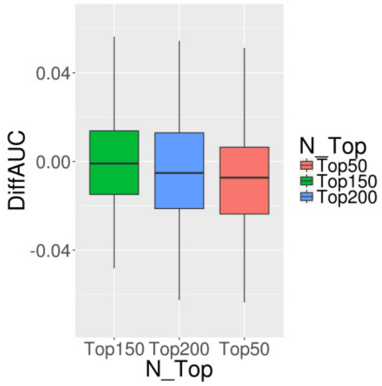

(a)

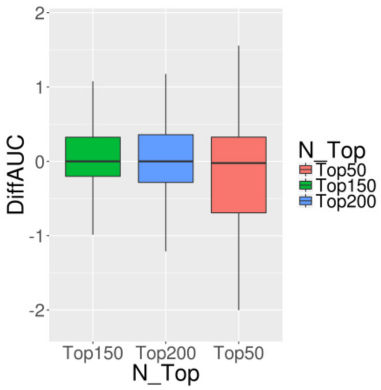

(b)

Figure S3. Distribution of differences in AUC and rpAUC of using 50, 150, and 200 signature genes compared to using 100 signature genes. (a) Difference in AUC. (b) Difference in rpAUC (FPR<0.1).

##### *F. Associating mTOR PAS with signatures of perturbations of MTOR pathway*

To assess the association between a perturbation signature and mTOR PAS, signature genes were divided into two groups based on the direction of their activity scores (positive vs negative). The association between the changes in expression after perturbation for the signature genes and the positive/negative grouping was assessed by the difference in the average TS values of the two groups (positive/negative) of signature genes. In Fig 2B and 2C, we assess the differences in associations between signatures of chemical perturbagens targeting proteins in the mTOR pathway and the control DMSO signatures. For the 5 CP signatures targeting AMPK, we flipped the signs of the differences in the average TS values to make them comparable with those of the other signatures. In Fig 2D we display the association with signatures of PI-103 treatment and the signature of amino acid starvation as the negative log of the p-value for the t-test comparing average expression changes between positive and negative signature genes.

##### *G. Creation of external mTOR pathway perturbation signatures*

Two microarray datasets profiling transcriptional responses of mTOR pathway perturbations (Guillard, et al., 2009; Tang, et al., 2015) were identified and downloaded from GEO. The first dataset (GSE12175(Guillard, et al., 2009) ) contained two-channel microarray data profiling the response of two glioma cell lines to treatment by the dual PI3K/mTOR inhibitor, PI-103. Raw GenePix® files (.gpr) from a two-color spotted cDNA microarray were downloaded from GEO. The raw data were corrected for background intensity and quantile-normalized using the *limma* R package(Ritchie, et al., 2015). PI-103 treated samples were compared to untreated samples using the empirical Bayes linear model as implemented in the *limma* R package to create differential expression profiles. The second dataset (GSE62673) contained Affymetrix® microarray expression data profiling the response of the MCF7 cell line starved of amino acids(Tang, et al., 2015). The raw data were RMA-normalized(Irizarry, et al., 2003) using the *affy* R package. Differential expression profiles between treated (amino acid starved) and untreated (full amino acid) samples were created using empirical Bayes linear model as implemented in the *limma* R package.

##### *H. Node score results of S6K across cell lines*

We applied our node score calculation to S6K1 and S6K2 proteins respectively to the 12 cell lines individually. The significance of positive contribution of a node is assessed by the Wilcoxon signed rank test. The cell lines HA1E, HT29, MCF7, and PC3 are found to have significantly positive contribution when both S6K proteins are located as inhibitors of IRS1, respectively.

| | Cell | S6K1 | | | S6K2 | | |
|---|---|---|---|---|---|---|---|
| | | Mean | Median | p.value | Mean | Median | p.value |
| 1 | A375 | 0.023 | -0.010 | 0.3936 | 0.028 | 0.009 | 0.2110 |
| 2 | A549 | 0.089 | 0.081 | 0.0000 | NA | NA | NA |
| 3 | HA1E | 0.099 | 0.110 | 0.0000 | 0.069 | 0.082 | 0.0000 |
| 4 | HCC515 | NA | NA | NA | NA | NA | NA |
| 5 | HEPG2 | -0.011 | 0.000 | 0.6976 | 0.162 | 0.176 | 0.0000 |
| 6 | HT29 | 0.119 | 0.118 | 0.0000 | 0.040 | 0.068 | 0.0098 |
| 7 | MCF7 | 0.182 | 0.138 | 0.0000 | 0.133 | 0.140 | 0.0000 |
| 8 | NPC | 0.039 | 0.028 | 0.0008 | NA | NA | NA |
| 9 | PC3 | 0.083 | 0.110 | 0.0000 | 0.079 | 0.125 | 0.0000 |
| 10 | SW480 | NA | NA | NA | NA | NA | NA |
| 11 | VCAP | 0.107 | 0.097 | 0.0000 | -0.011 | -0.010 | 0.8211 |
| 12 | HEKTE | NA | NA | NA | NA | NA | NA |

Table S3. Node score of S6K when located as an inhibitor of IRS1. “NA” indicates the KD signature of the corresponding node is not available.

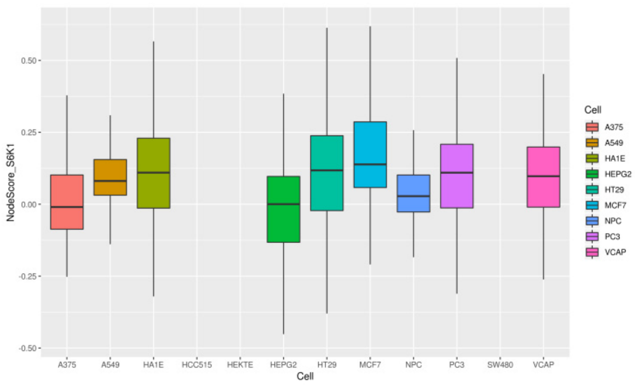

(a) S6K1

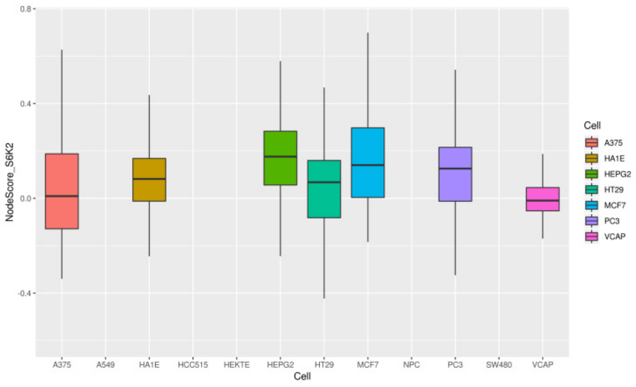

(b) S6K2

Figure S4. Distribution of node score of S6K across 12 cell lines. Scores are calculated with locating the node as an inhibitor of IRS1.

#### *I. Figure S5: Heatmap of AUC and rpAUC for the comparison of four methods*

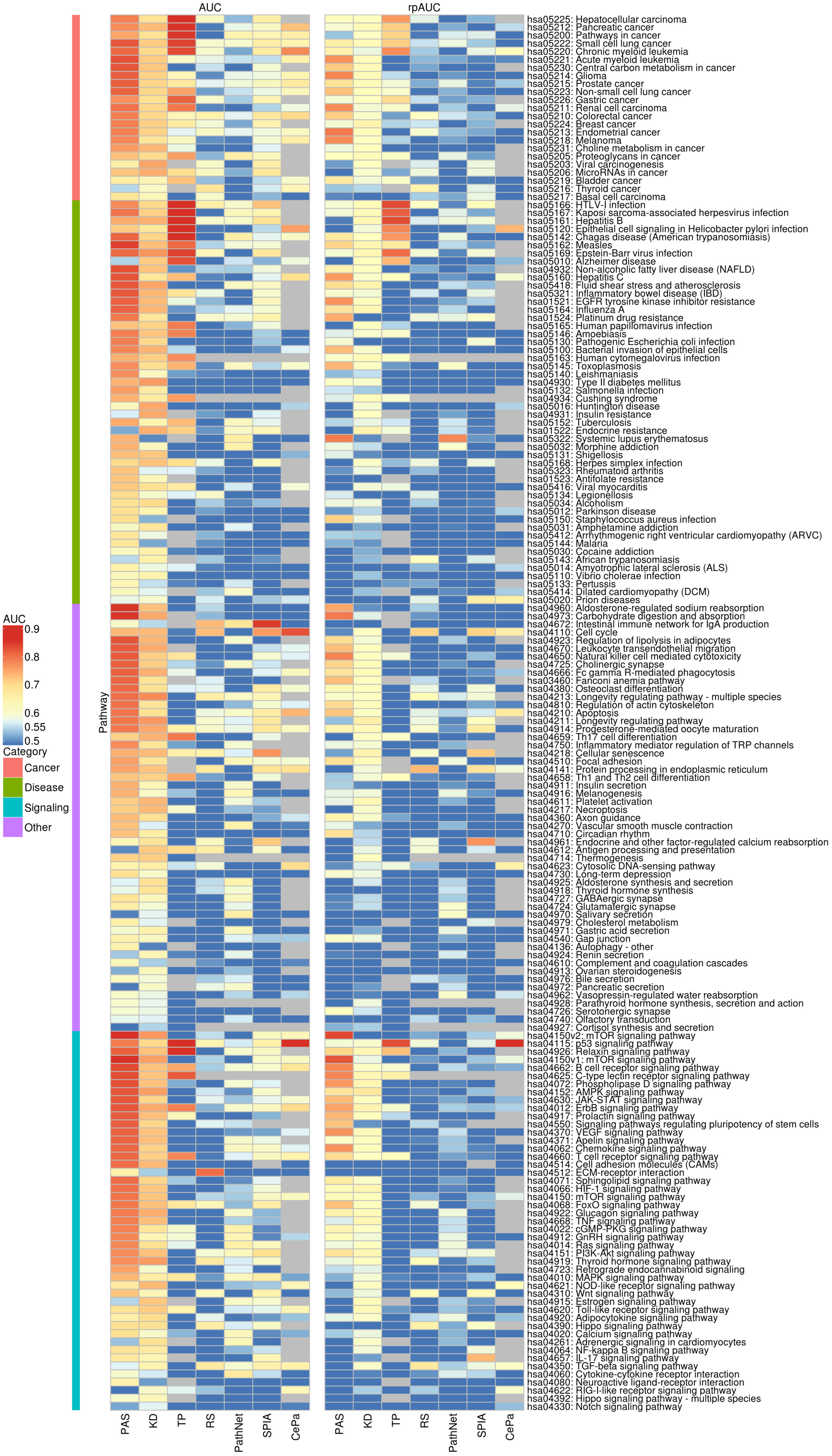

***J. Figure S6: Summary results for rpAUC comparisons of seven methods***

Percentage of pathways predicted with the highest AUC for seven different methods across four different types of KEGG pathways; C) Heatmap of AUC's for predicting affected KEGG signaling pathways for all seven methods.

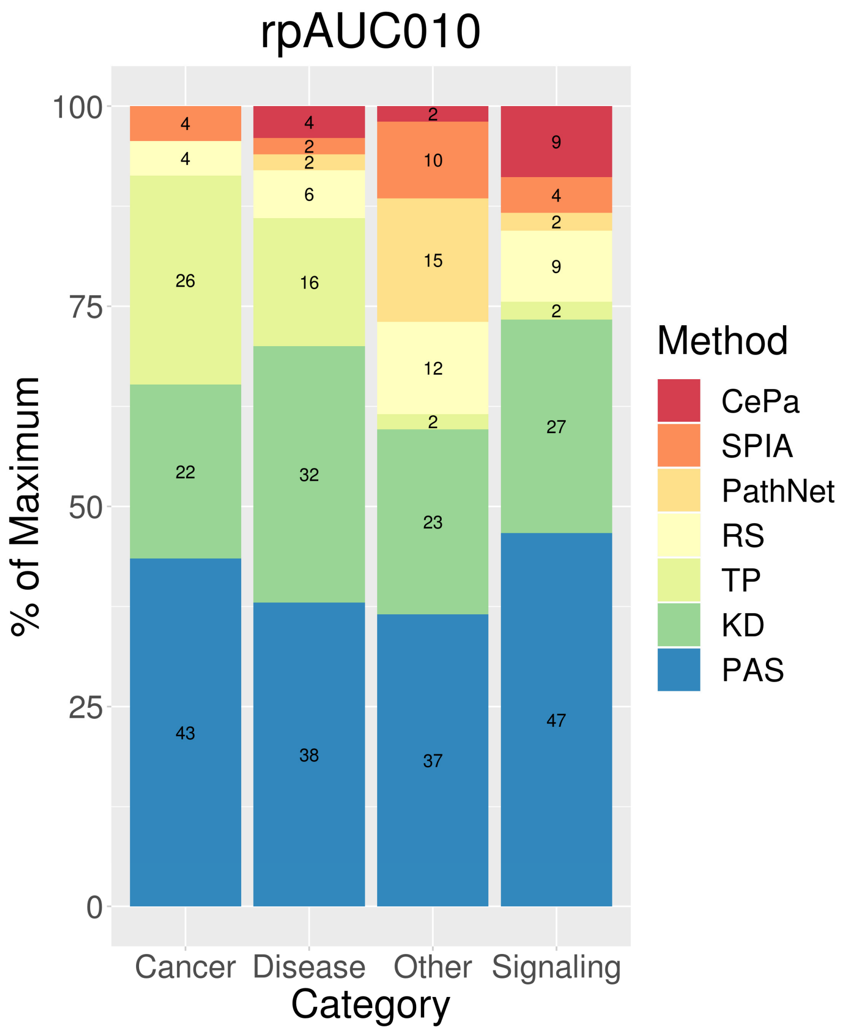
